## Supplementary figures and images for "Basigin mediation of *Plasmodium falciparum* red blood cell invasion does not require its interaction with monocarboxylate transporter 1"

### Supplementary Figure 1

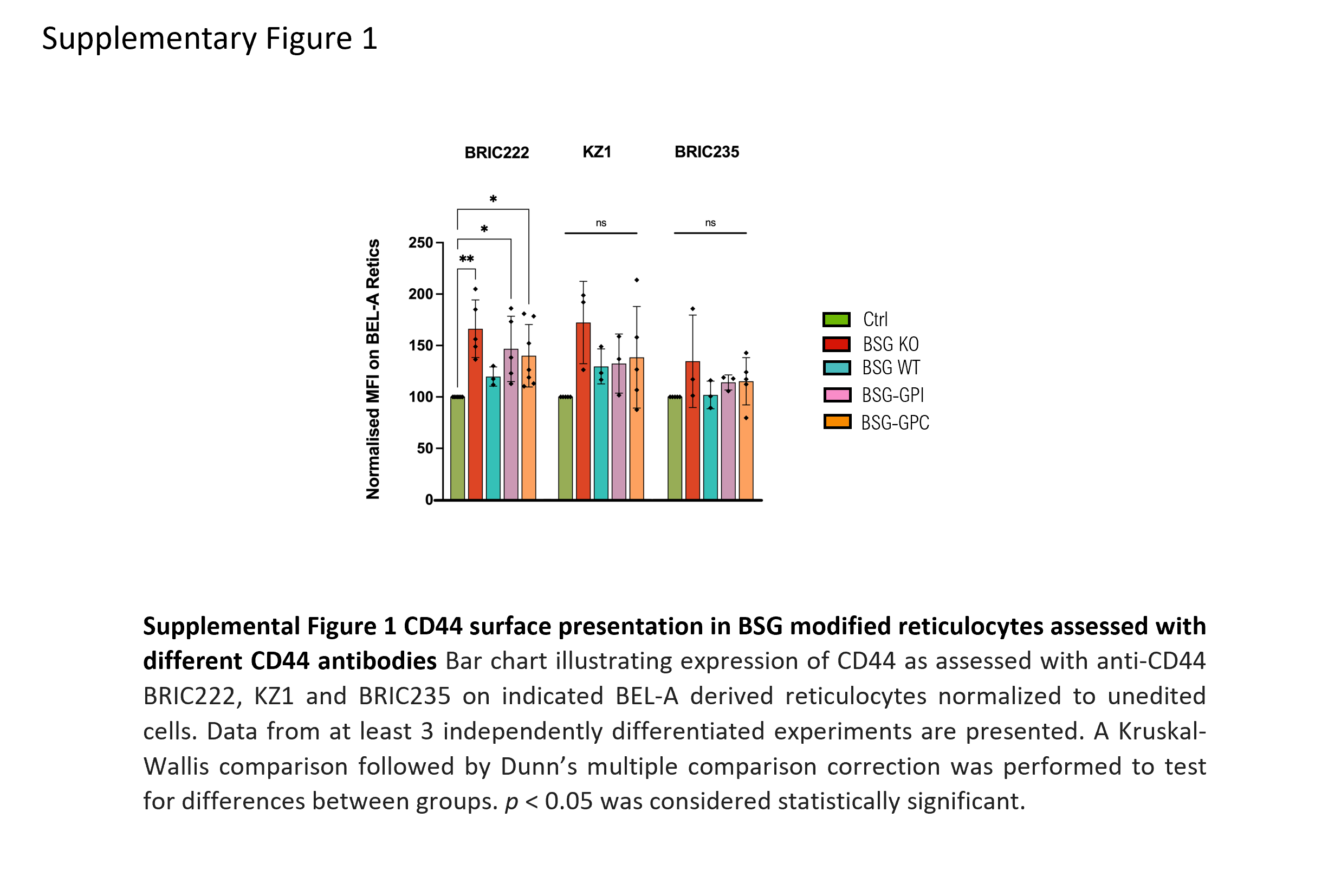
